# Differential labelling of human sub-cellular compartments with fluorescent dye esters and expansion microscopy

**DOI:** 10.1101/2023.02.21.529394

**Authors:** Thomas M. D. Sheard, Tayla Shakespeare, Rajpinder S. Seehra, Michael E Spencer, Kin M. Suen, Izzy Jayasinghe

## Abstract

Amine-reactive esters of aromatic fluorescent dyes are emerging as imaging probes for nondescript staining of cellular and tissue architectures. We characterised the differential staining patterns of 14 fluorescent dye ester species with varying physical and spectral properties in the broadly studied human cell line – HeLa. When combined with expansion microscopy (ExM), these stains reveal nanoscale features such as the nuclear proteome, membrane-bound compartments and vesicles. Among N-Hydroxysuccinimide (NHS) esters, we observe differential compartment specificity and weighting of labelling density which correlates with the hydrophobicity of the dye ester. We also observe changes in both staining density and compartment specificity for a given dye ester depending on the presence of a second dye ester species and on the timepoint of application in the ExM protocol. Our findings confirm these dye esters as a useful addition to the repertoire of biomedical stains of the cellular proteome, either on their own, or as counterstains to immunofluorescence.

## Introduction

As the most broadly used analytical approach, modern microscopy techniques offer a view into the finest of length-scales in which the fundamental mechanisms of life operate. Variants of electron microscopy (EM) and super-resolution optical microscopies have been the primary tools for visualising the nanometre-scale features of biomolecular assemblies, cytoskeletons, membrane-bound, and membraneless compartments in cells. Due to the lack of intrinsic contrast in most biological samples, both approaches leverage staining techniques that are either passive (e.g. heavy metal, uranium and lead-based stains for EM) or targeted stains. The latter is commonly employed in fluorescence optical microscopies, with optimum specificities achieved with immuno-labelling and click chemistry strategies. Super-resolution techniques in particular rely on the contrast and specificity offered by targeted fluorescent probes.

A recent technique called expansion microscopy (ExM) achieves improved resolution by physically expanding an imprint of the sample inside a swellable acrylamide hydrogel which can then be imaged at super resolution with straightforward optical microscopy (Chen et al., 2015). ExM also involves proteolytic digestion of the sample, allowing it to expand physically. This effectively ‘decrowds’ its intrinsic molecular structures, significantly reducing levels of background signal to enhance the targets of interest. While fluorescence microscopy approaches like ExM have enabled extraordinary insights into the distribution of molecular targets, it has proven less useful for illustrating broader cellular architecture.

A novel labelling strategy to visualise cellular compartments entails the re-purposing of amine reactive N-Hydroxysuccinimide (NHS) esters, used commonly for fluorescently tagging purified proteins such as antibodies. When directly applied to samples containing cells, tissues or whole organisms, these fluorescent esters produce a nondescript staining of the entire cellular proteome and differentiation between organelles, cell types and tissue types. NHS esters have recently been introduced in pan-ExM (M’Saad & Bewersdorf, 2020), capitalising on the decrowding effect from 16-fold expansion to observe the finest of cellular ultrastructures such as mitochondrial cristae, golgi cisternae and nucleolar sub-compartments. Several other groups have used NHS esters with ExM (Damstra et al., 2022; Lee et al., 2022; Louvel et al., 2022; Mao et al., 2020; Sim et al., 2022), with many others implementing NHS esters alongside specific protein labels in order to better understand the biology of a range of cell systems (Bertiaux et al., 2021; Chacko et al., 2023; Hinterndorfer et al., 2022; Laporte et al., 2022; M’Saad et al., 2022; Rashpa & Brochet, 2022; Suen et al., 2022; Yu et al., 2020). The same probes and method have enabled unprecedented visualisations of the conformations of individual proteins, as shown with one nanometer ExM (Shaib et al., 2022).

While the rapid uptake of fluorescent NHS esters in the ExM community shows the promise of these labels, a number of key parameters still limits their reproducibility, validation and broader uptake within the bioimaging community. Firstly, the physical properties of the various dye ester species, in particular the aromatic dye molecule, have been hypothesised as the key determinants of the density of staining of a given compartment or cellular structure (M’Saad et al., 2022; Sim et al., 2022). For example, hydrophobic ester species may be attracted to lipid or polymer-rich compartments of the cell. However, the grammar of this differential staining remains poorly understood. Whilst antibody-based, targeted co-staining promises to be a useful strategy to establish this grammar, interactions of the nondescript stains with the targeted probe itself and the consequences to the heterogeneous labelling densities observed in the super-resolution image remain poorly understood. Only a limited number of dye ester species have been tested so far; hence, there remains to be a more systematic evaluation of a broad range of fluorescent dye esters.

In this report, we characterised a variety of fluorescent dye ester species with wide-ranging physical and spectral properties to label different cellular compartments in HeLa cells. We demonstrated distinct compartment labelling between different ester dye families, with ExM, and benchmark alongside antibodies. Presented below is a series of complexities and time-dependencies that we have systematically observed in nondescript (often multiplexed) staining with NHS esters that pose considerable implications for using them as stains for imaging human cells.

## Results

### Ester catalogue characterisation

We characterised a selection of 13 fluorescent dye ester species belonging to different dye families (Alexa Fluor, AZ, BODIPY, ATTO, and MB), each featuring different charges, hydrophobic properties and spectral range of fluorescence emission (Figure 1A). The hydrophobicity of each of these molecules relates to how nonpolar groups behave in an aqueous environment, and is predicted from the distribution coefficient logD value (listed in Table 1), calculated using their structural formula. By this characterisation, dye ester species with more negative logD values are more hydrophilic; dyes with more positive logD values are hydrophobic.

**Figure 1.**
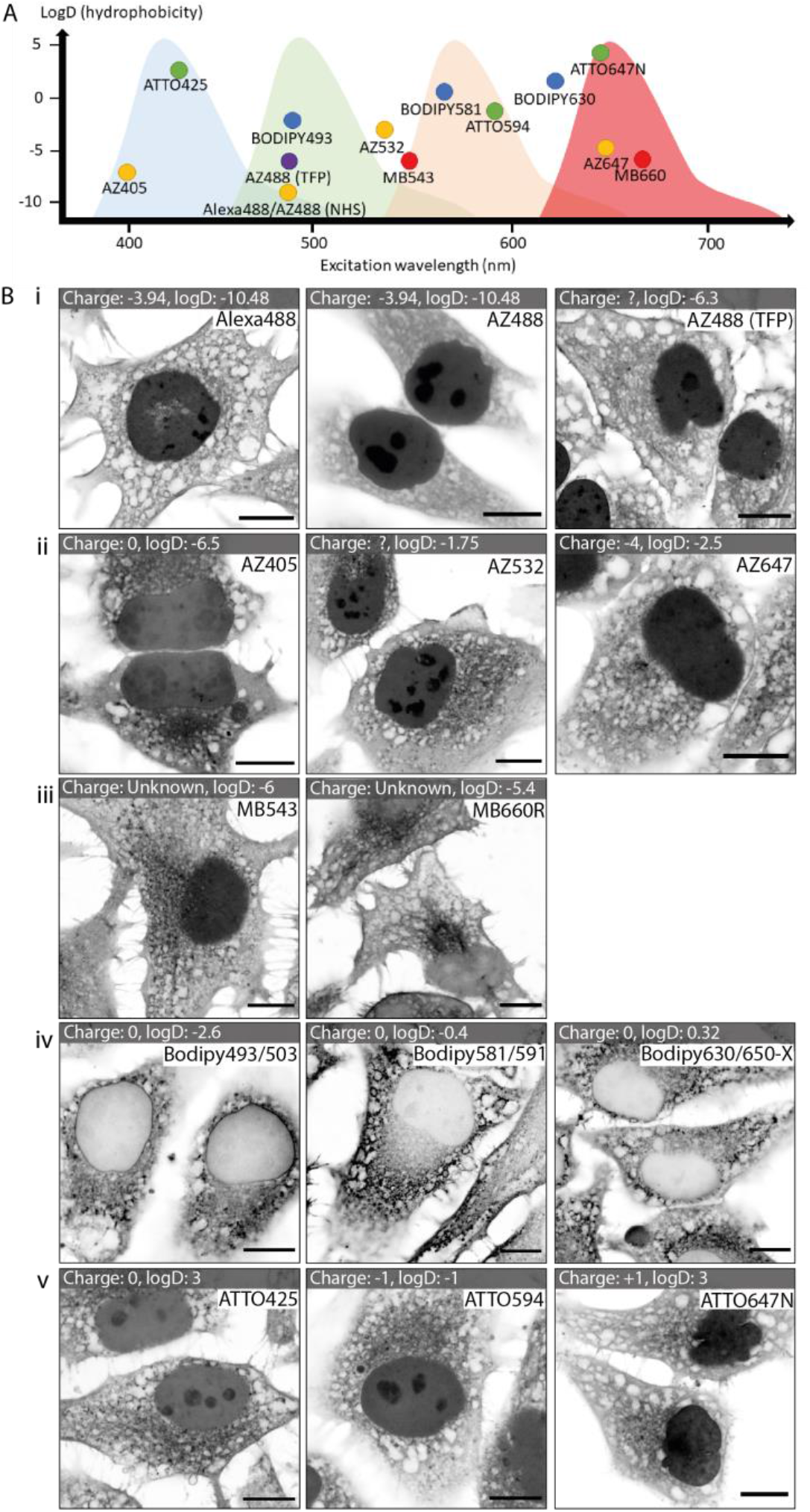
Catalogue of dye ester images. **(A)** Chart displaying the ester dyes, distributed based on their excitation wavelength and hydrophobicity (determined from logD values). **(B)** A gallery of unexpanded images of each of the esters. Between different dye families there are substantial differences, and within a single dye family there are similarities in patterns. Scale bars: 10 μm.

**Table 1.** Ester properties. Details of each ester-dye conjugate are provided, specifically the light maximum excitation and emission wavelengths, and the chemical properties of hydrophobicity (inferred from the logD value, D being the ratio of the solute between a nonpolar and a polar solvent) and the charge. The more positive value of logD, the more hydrophobic the molecule. Information was obtained from the following sources: ^a^(Zanetti-Domingues et al., 2013), ^b^(Hughes et al., 2014), ^c^(Zhang et al., 2017), ^d^(URL: https://www.spectra.arizona.edu/supplemental/ATTO_Dye_Properties_01.pdf, accessed 15/02/2023), *(calculated using Chemaxon logD predictor).

| Dye | Exc. Max (nm) | Em. Max (nm) | Hydrophobicity (logD) | Overall Charge | Source |
| --- | --- | --- | --- | --- | --- |
| NHS Alexa488 | 494 | 517 | (-10.48)<sup>a</sup> | (-3.94)<sup>a</sup> | Thermo Fisher Scientific |
| NHS AZ488 | 490 | 525 | (-9.4)* | (-3.94)<sup>a</sup> | Fluoroprobes |
| TFP AZ488 | 495 | 515 | (-6.3)* | Unknown | Fluoroprobes |
| NHS AZ405 | 401 | 421 | (-6.5)* | Unknown | Fluoroprobes |
| NHS AZ532 | 532 | 554 | (-3.26)<sup>b</sup>, (-1.75)* | Unknown | Fluoroprobes |
| NHS AZ647 | 650 | 665 | (-6.72)<sup>b</sup>, (-2.5)* | (-4)<sup>c</sup> | Fluoroprobes |
| NHS BODIPY493/503 | 493 | 503 | (-2.6)* | 0 | Thermo Fisher Scientific |
| NHS BODIPY581/591 | 581 | 591 | (-0.4)* | 0 | Thermo Fisher Scientific |
| NHS BODIPY630/650-X | 625 | 640 | (0.32)* | 0 | Thermo Fisher Scientific |
| NHS ATTO425 | 439 | 485 | (3)* | (0)<sup>d</sup> | Sigma-Aldrich |
| NHS ATTO594 | 597 | 625 | (-1)* | (-1)<sup>d</sup> | Sigma-Aldrich |
| NHS ATTO647N | 645 | 669 | (1.96)<sup>a</sup>, (3)* | (1)<sup>d</sup>, (0.61)<sup>a</sup> | Sigma-Aldrich |
| NHS MB543 | 543 | 566 | (-6)* | 1 more negative sulfo group vs equivalent Alexa dye | Fluoroprobes |
| NHS MB660R | 673 | 694 | (-5.4)* | 1 more negative sulfo group vs equivalent Alexa dye | Fluoroprobes |

**Table 2.** Key Resources. Details of cell lines, reagents, antibodies, esters and softwares are provided.

|  | SOURCE | IDENTIFIER |
| --- | --- | --- |
| Experimental models: Cell lines |  |  |
| Human: HeLa cell | Sigma | Cat# 93021013 |
| Chemicals |  |  |
| Dulbecco's Modified Eagle Medium | Thermo Fisher Scientific | Cat# 11584496 |
| Foetal bovine serum | Labtech | Cat# SKU: FB-1001/100-500 |
| Penicillin-streptomycin | Thermo Fisher Scientific | Cat# 11548876 |
| Poly-D-lysine | Cultrex | Cat# 3439-100-01 |
| Paraformaldehyde | Sigma-Aldrich | Cat# P6148-500G |
| Bovine serum albumin | Thermo Fisher Scientific | Cat# 30063-572 |
| Sodium azide | Sigma-Aldrich | Cat# S8032-25G |
| DMSO | Biotium | Cat# 90082-BT |
| Triton X-100 | Sigma-Aldrich | Cat# T9284-100ML |
| Normal goat serum | Thermo Fisher Scientific | Cat# 10000C |
| Acryloyl-X | Thermo Fisher Scientific | Cat# A20770 |
| Acrylamide | Sigma-Aldrich | Cat# A9099-25G |
| N,N'-Methylenebisacrylamide | Sigma-Aldrich | Cat# M7279-25G |
| Ammonium persulfate | Sigma-Aldrich | Cat# A3678-25G |
| N,N,N',N'-Tetramethylethylenediamine | Sigma-Aldrich | Cat# T7024-25ML |
| Proteinase K | New England Biolabs | Cat# P8107S |
| Ethylenediaminetetraacetic acid | Sigma-Aldrich | Cat# EDS-100G |
| Guanidine HCl | Sigma-Aldrich | Cat# G3272-25G |
| Tris pH 8.0 | Invitrogen | Cat# P8920-100ML |
| Poly-L-lysine | Sigma-Aldrich | Cat# 10259194 |
| Dye-esters |  |  |
| NHS Alexa488 | Thermo Fisher Scientific | Cat# a20000 |
| NHS AZ488 | Fluoroprobes | Cat# 1013-1 |
| TFP AZ488 | Fluoroprobes | Cat# 1026 |
| NHS AZ405 | Fluoroprobes | Cat# 1061-1 |
| NHS AZ532 | Fluoroprobes | Cat# 1041-1 |
| NHS AZ647 | Fluoroprobes | Cat# 1121-1 |
| NHS BODIPY493/503 | Thermo Fisher Scientific | Cat# D2191 |
| NHS BODIPY581/591 | Thermo Fisher Scientific | Cat# D2228 |
| NHS BODIPY630/650-X | Thermo Fisher Scientific | Cat# D10000 |
| NHS ATTO425 | Sigma-Aldrich | Cat# 16805 |
| NHS ATTO594 | Sigma-Aldrich | Cat# 8741 |
| NHS ATTO647N | Sigma-Aldrich | Cat# 18373 |
| NHS MB543 | Fluoroprobes | Cat# 1661-1 |
| NHS MB660R | Fluoroprobes | Cat# 1661-1 |
| Antibodies |  |  |
| Rabbit polyclonal KDEL | Thermo Fisher Scientific | Cat# PA1-013 |
| Rabbit monoclonal GM130 | Abcam | Cat# ab52649 |
| Mouse monoclonal ATP5A1 | Thermo Fisher Scientific | Cat# 43-9800 |
| Alexa Fluor 488 goat anti-mouse IgG | Thermo Fisher Scientific | Cat# A11001 |
| Alexa Fluor 488 goat anti-rabbit IgG | Thermo Fisher Scientific | Cat# A11008 |
| Alexa Fluor 594 goat anti-mouse IgG | Thermo Fisher Scientific | Cat# A11005 |
| Alexa Fluor 594 goat anti-rabbit IgG | Thermo Fisher Scientific | Cat# A11012 |
| Atto 647N goat anti-rabbit IgG | Sigma-Aldrich | Cat# 40839-1ML-F |
| Atto 647N goat anti-mouse IgG | Sigma-Aldrich | Cat# 50185-1ML-F |
| Items |  |  |
| Glass coverslips, #1.5, 22x22 mm | Menzel-Glaser | Cat# 631-0851 |
| Chamberslides | Razorlab | Custom-made |
| Software and algorithms |  |  |
| Fiji | ImageJ | https://imagej.net/software/fiji/ |
| Chemaxon calculator plugin | Chemaxon | https://disco.chemaxon.com/calculators/demo/plugins/logd/ |
| Excel | Microsoft |  |

A catalogue of standard Airyscan images shows the labelling patterns for each ester in fixed, unexpanded HeLa cells (Figure 1B). The Alexa dyes (and their derivatives) generally feature a negative charge due to sulfonation, and are thus more hydrophilic (Panchuk-Voloshina et al., 1999). NHS Alexa488 labelled a web-like pattern that resembled the endoplasmic reticulum (ER), as well as within the nucleus (Fig 1B-i). The AZ family of dyes are structurally identical to the Alexa derivatives, however are available at a cheaper cost through a different manufacturer. Consequently, NHS AZ488 labelling pattern shared strong qualitative similarities to that of NHS Alexa488. The tetrafluorophenyl (TFP) ester of AZ488, despite being more stable in the linkages it forms in comparison to the equivalent NHS ester, is less hydrophilic (see Table 1). By comparison with Alexa 488 and AZ488 NHS ester, the TFP ester staining appeared more intricate, likely as a result of staining more lipid dense compartments or vesicles in addition to ER (see comparison in Fig 1B-i). Other AZ dyes that we tested included AZ405, AZ532, AZ647 (Fig 1B-ii). These dye ester species all shared a mixture of labelling patterns across the cytoplasm and nucleus. The MB dye esters, also derived from Alexa dyes, feature an additional negatively charged sulfo-group (improving water solubility and minimising self-quenching) and are predicted to be similarly hydrophilic. MB543 appeared to stain cytosolic granules or vesicles with greater labelling density than the ER network, while MB660 more densely labelled a compartment adjacent to the nucleus, likely the golgi apparatus or rough ER (Fig 1B-iii).

In contrast to the hydrophilic Alexa and AZ dyes, the BODIPY family of dyes are strongly hydrophobic. Labelling of these dyes (BODIPY493/503, BODIPY581/591 and BODIPY 630/650-X) similarly included ER networks, however with an absence of nuclear labelling (Figure 1B-iv). A clear gradation of staining intensity was observed with intense staining of the nuclear envelope and perinuclear ER network (i.e. rough ER, rich in integral membrane proteins consisting of lipophilic domains), and diminishing intensity near the cellular extremities. The ATTO family of dyes typically features more positive logD values (Table 1) than Alexa dyes, however can exhibit variable water solubilities (Patra et al., 2020). This variability was reflected in the HeLa cell labelling patterns across NHS ATTO425, ATTO594 and ATTO647N that stained nuclei with differing densities, also reporting ER and perinuclear compartments and varying staining densities (Figure 1B-v).

### Differential compartment labelling with fluorescent dye esters

We used the 4x enhanced ExM (EExM) approach, detailed previously (Sheard et al., 2019; Sheard & Jayasinghe, 2020). This approach allowed us to compound the 4-fold resolution improvement with ExM with the resolution-doubling of Airyscan microscopy to achieve an effective resolution of ~ 40 nm. The hydrophobic dye ester NHS BODIPY493/503 (with a net charge of 0) yielded an impression of a variety of cellular compartments, for instance vesicular shapes resembling lipid droplets, peroxisomes and mitochondria (Figure 2A, left). The detail of compartment labelling is qualitatively comparable to the equivalent electron micrographs published by Hennies et al. (Hennies et al., 2020) (Figure 2A, right), albeit electron microscopy still offers greater contrast owing to thin sectioning and superior resolution.

**Figure 2.**
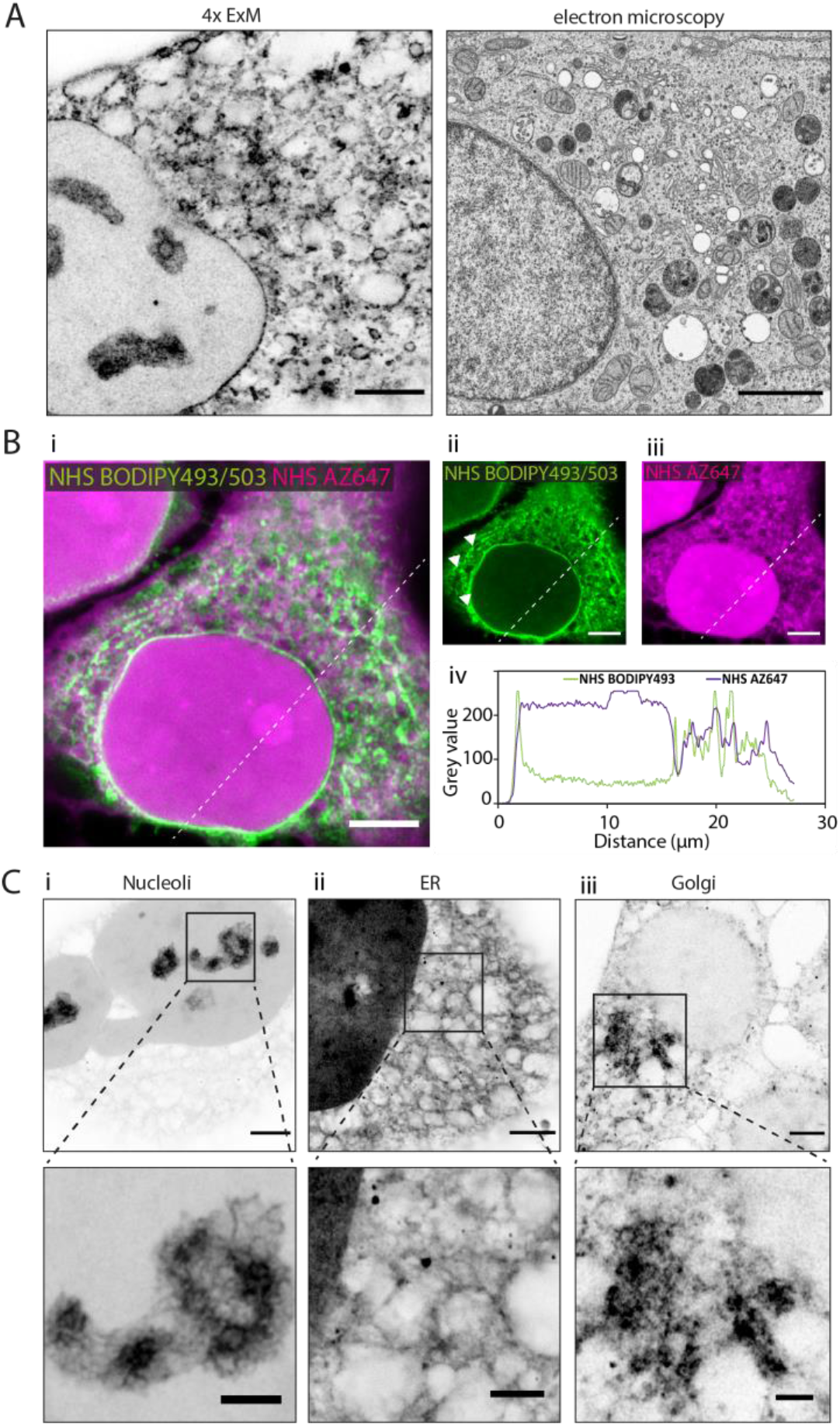
Distinct compartments visualised with NHS esters and expansion microscopy. Esters enable mapping of cellular compartments when conjugated with fluorescent dyes. **(A)** Intracellular compartments can be observed in a HeLa cell labelled with NHS BODIPY493/503 and imaged with 4x EExM (left), revealing an equivalent level of spatial detail and distinction between cellular sub-compartments revealed in 2D transmission electron micrograph (right). Image adapted from Hennies et al. with permission (Hennies et al., 2020). **(B)** Preexpansion images show how the ester labelling pattern differs depending on the properties of the conjugated dye. **(i-iii)** The hydrophilic NHS AZ647 (magenta) labels the nucleus preferentially than the hydrophobic NHS BODIPY493/503 (green) as indicated by the increased intensity shown in the line profile chart **(iv)**. **(iv)** Modest, positive correlation in the non-nuclear regions and compartments, indicated by the line intensity profile. **(C)** 4x EExM images demonstrate some of the sub-cellular compartments that can be preferentially labelled by esters, including **(i)** nucleoli (NHS MB543 applied pre-gelation), **(ii)** the membranous web of endoplasmic reticulum (NHS AZ532 applied pre-gelation), and **(iii)** dense cytoplasmic regions adjacent to the nucleus belonging to the golgi apparatus (NHS BODIPY630 applied post-digestion). Scale bars (expansion factor rescaled): (A) (left) 2.5 μm, (right) 2 μm; (B) 10 μm, (C) (upper) 2.5 μm, (lower) 4 μm.

We directly compared the labelling patterns from dye ester species with opposing physical properties. A comparison of standard Airyscan images of a HeLa cell staining (Fig 2B-i) with hydrophobic NHS BODIPY493/503 (green, ii) and the hydrophilic NHS AZ647 (magenta, 5 iii) shows that differential, dual staining is achievable with spectrally distinct dye ester species. Both esters yielded weakly correlated labelling of non-nuclear compartments (Fig 2B-iv). With NHS BODIPY493/503, there were more elongated structures visible surrounding the nucleus, resembling mitochondria (arrowheads in panel B-ii). The most striking differences related to the differential staining of the nuclear proteome with NHS AZ647 staining strongly and a distinct absence of NHS BODIPY493/503 from the nucleoplasm – difference that is attributable to the differing hydrophobic and charge profiles of the dyes. By contrast, the highly lipophilic BODIPY493/503 preferentially labelled nuclear envelopes and rough ER, likely containing high densities of integral membrane proteins. Combining these stains with 4x EExM (i.e. the integration of 4x ExM with Airyscan imaging) revealed highly detailed images of nucleoli and the chromatin associated proteome (Fig 2C-i, NHS MB543 ester), rough ER featuring protein densities resembling ribosomes (Fig 2C-ii, NHS AZ532 ester), and perinuclear Golgi apparatus (Fig 2C-iii, NHS BODIPY630 ester).

#### Validation of the labelling patterns with dual antibody-ester labelling

Immunofluorescence labelling experiments often have the drawback of lacking an ultrastructural context against which the targets localised by the antibodies. We investigated the compartment specificities of dye esters (applied pre-gelation) through triple multiplexed imaging of Golgi apparatus (via anti-GM130; yellow) and mitochondria (via ATP synthase ATP5A1; magenta) against a counterstain of Alexa488 NHS ester stain (cyan), shown in in Figure 3A. Whilst the perinuclear staining with the Alexa488 NHS ester resembled a web-like network, this result revealed that the structure observed is a composite of the geometries formed by multiple compartments, including mitochondria and Golgi apparatus. It was nevertheless useful for differentiating cellular spaces devoid of the major compartments and the boundary of the nucleoplasm. In a further comparison, antibody labelling for KDEL (a common target peptide sequence encoding for their retention or returning to the ER) revealed geometries of the ER network within the full cytoplasmic architecture, as displayed within NHS AZ488 labelling (magenta) (Figure 3B-i). The KDEL labelling was weighted heavily on smooth ER, while the dye ester label traced a more complete overall impression of ER network organisation. The mitochondria were visualised with antibody labelling of ATP5A1 with coinciding counter staining with NHS ATTO647N (Figure 3Bii). Finally, the cis-golgi has been visualised by antibody labelling for GM130, and appeared to be preferentially labelled by the hydrophobic NHS BODIPY493/503, which also reported the closely positioned nuclear membrane (Figure 3Biii).

**Figure 3.**
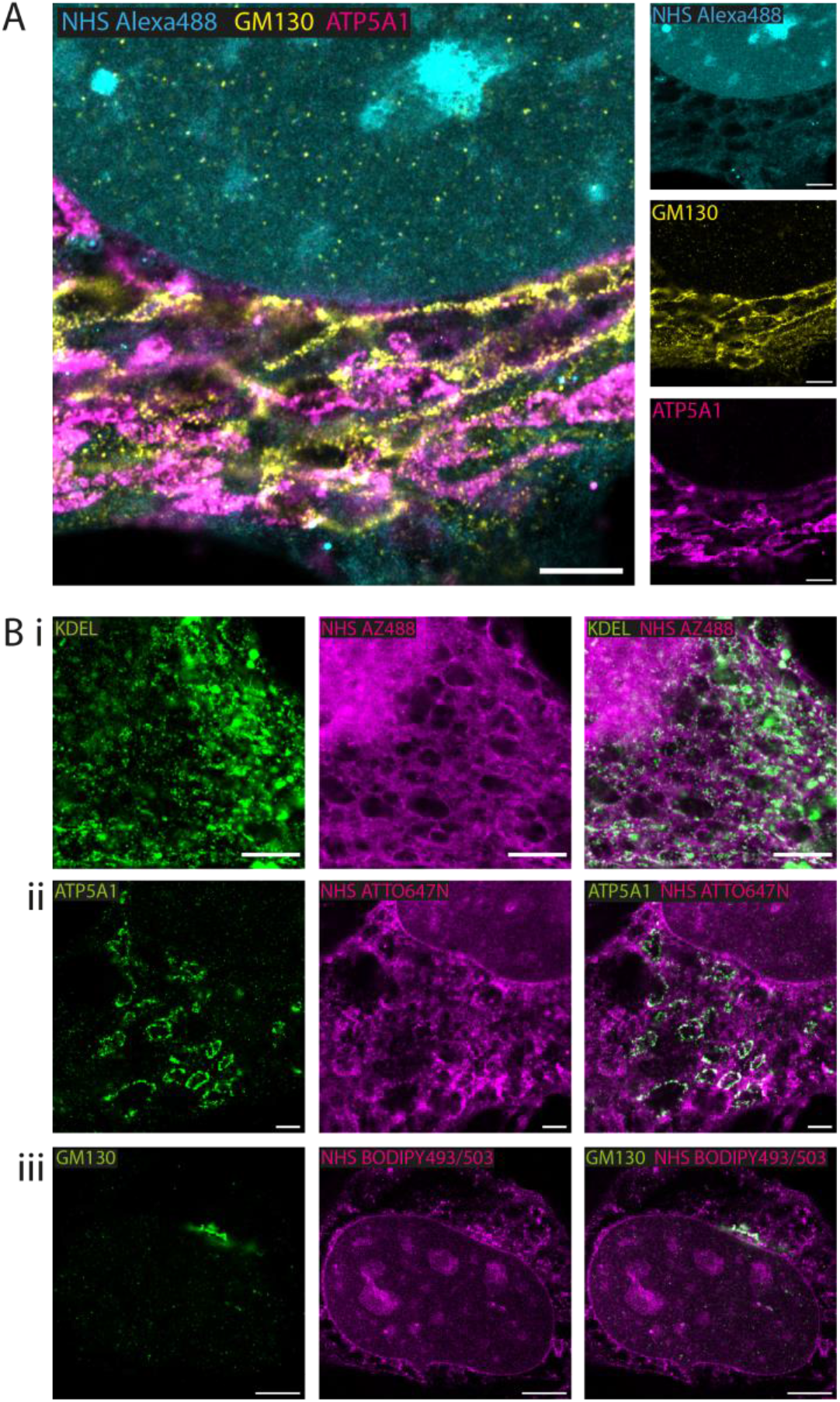
Visualising compartments alongside antibody benchmarks. By implementing NHS esters alongside antibodies it is possible to get context for those specific protein targets in relation to the broader cellular architecture. **(A)** 4x EExM images of NHS Alexa488 (cyan) alongside antibody labels for cis-golgi marker GM130 (yellow) and the mitochondrial ATP5A1 ATP synthase (magenta). **(B)** Gallery of 4x EExM images showing examples of the preferential compartment labelling by esters (all applied pre-gelation) alongside immunolabelled compartments, including **(i)** the endoplasmic reticulum network (as labelled with antibodies recognising KDEL in the lumen) alongside NHS AZ488, **(ii)** the mitochondria as shown by immunolabelling for ATP5A1, alongside NHS ATTO647N, and **(iii)** the golgi, targeted with GM130 antibodies, alongside NHS BODIPY493/503. Scale bars (expansion factor rescaled): (A) 3.75 μm; (Bi) 2.5 μm, (Bii) 2.5 μm, (Biii) 5 μm.

### Establishing the grammar of staining HeLa cells with fluorescent NHS esters

#### Application of dye ester at different time-points of the ExM protocol impacts labelling specificity

The sequence of application of the fluorescent NHS ester species has been hypothesised to impact the labelling densities and the cellular compartments that they report with ExM (Sim et al., 2022). It has been hypothesised that the staining patterns of the dye ester is impacted by the presence of the hydrogel and the ‘decrowding’ imparted by the proteolytic digestion and expansion. The specificity of the dye ester as an imaging probe could therefore be impacted by the stage of the ExM protocol at which it is applied. To investigate this, we applied NHS ATTO647N at three time-points: pre-gelation, inter-digestion (4 hours digestion, labelling, followed by 4 hours further digestion), and at post-digestion within separate samples (Supplementary Figure 1A). When applied pre-gelation NHS ATTO647N stained elongated compartments surrounding the nuclei that strongly resembled mitochondria. A moderate density of staining was also present within the nucleoplasm (Supplementary Figure 1B). In contrast, when applied in an inter-digestion window the labelling was completely excluded from the nucleoplasm, and the cytoplasmic labelling appeared to concentrate at discrete protein densities (resembling either rough ER or small vesicles). The labelling pattern observed in the pre-gelation stage was completely reversed when the same dye ester was applied post-digestion stage, most strongly labelling the nucleus, while the cytoplasmic labelling appears weaker in intensity and the protein densities sparser.

#### Competitive interactions between dye ester species

We sought to examine whether simultaneous application of NHS dye esters could shift their target specificity. We co-stained HeLa cells with hydrophilic TFP AZ488 at the same time as one of two esters of distinctly different logD values. In the first instance we paired TFP AZ488 (predicted logD of −6.3; see Table 1) with another hydrophilic dye - NHS AZ647 (predicted logD of −2.5). In standard Airyscan images of ester-labelled cells, the intensity of TFP AZ488 staining of the ER and non-nuclear compartments was strong (Fig 4A-i). In another experiment, we applied TFP AZ488 ester and the hydrophobic NHS ATTO647N (log D of +3) simultaneously. The TFP AZ488 labelling density the non-nuclear compartment in the latter experiment was considerably weaker in the presence of the second, hydrophobic dye (Fig 4A-ii). This difference in densities of labelling, particularly relative to that in the nucleoplasms, is shown in the comparative line profile plots (Figure 4A-iii).

**Figure 4.**
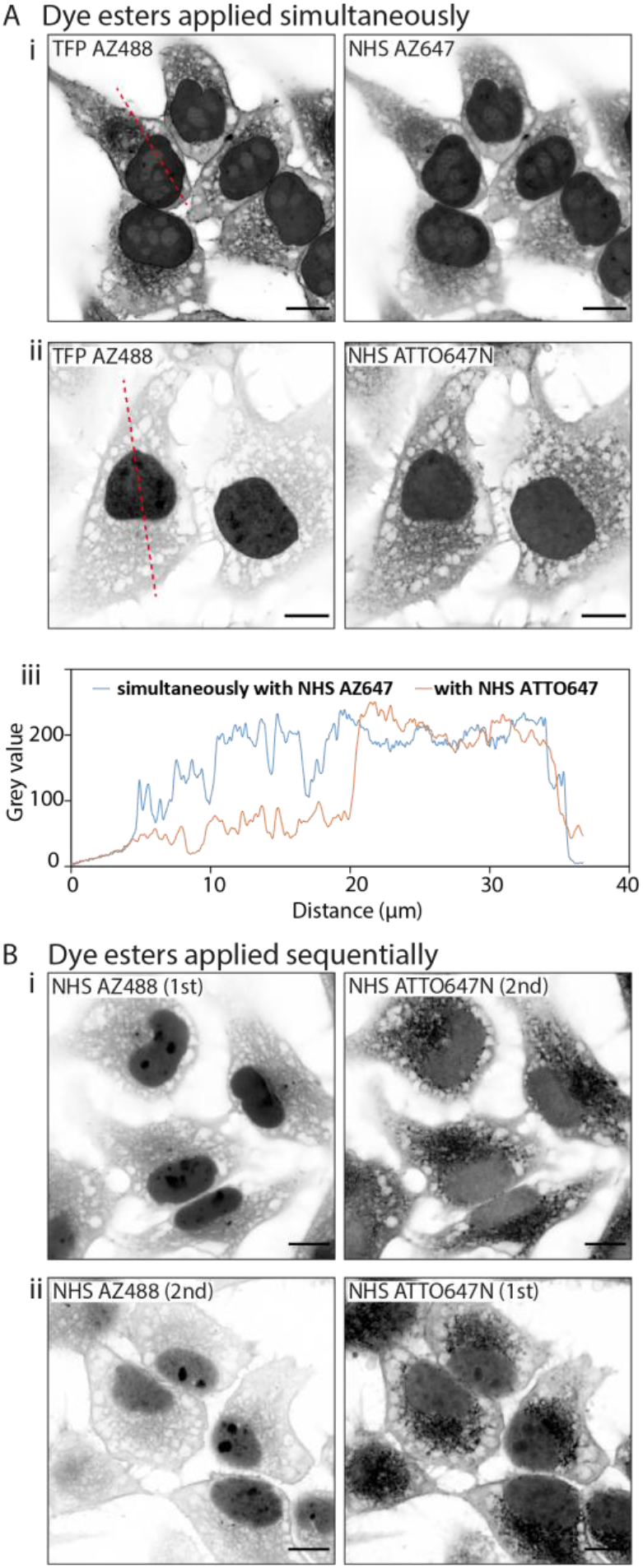
Interactions between multiple dye ester species. When multiple dye esters are added into a single sample, competitive interactions between ester species alter their labelling patterns. **(A)** Unexpanded images show that when esters are added simultaneously, (i) TFP AZ488 (which is hydrophilic) labels the cytoplasm and nucleus in a uniform way when paired with NHS AZ647 (also hydrophilic), (ii) but more strongly labels the nucleus when paired with NHS ATTO647N (hydrophobic). (iii) Line profile plot demonstrates the differing intensities across the cells. **(B)** When dye ester species are added sequentially their labelling can be altered. Shown, are examples of (i) applying NHS AZ488 first, then NHS ATTO647N second where the intensity of NHS AZ488 is much stronger than (ii) in the converse sequence. Scale bars: 10 μm.

#### Sequential application of dye ester species

We examined the implications of adding two esters sequentially by labelling cells firstly with hydrophilic NHS AZ488, and then with the hydrophobic NHS ATTO647N (Figure 4B-i). In the second experiment we switched the order around, first labelling with hydrophobic NHS ATTO647N followed by the hydrophilic NHS AZ488 (Figure 4B-i). In both situations the labelling patterns of NHS ATTO647N were qualitatively unchanged. However, when applied second the NHS AZ488 stained at a lower density, most notably in the nucleus (Figure 4B-ii).

## Discussion

### Up-take of esters in the microscopy community

The new approach of direct application of fluorescent dye esters as nondescript stains has dramatically expanded the repertoire of fluorescent labelling protocols for high- or super-resolution microscopies. The use of esters to label the proteome promises valuable contextual information for interpreting protein localisation and the overall structural phenotypes of the cell. With many different commercially available esters, a researcher can survey a few candidates and quickly discover labels allowing them an impression of the compartments of interest. The 14 dye ester species characterised in this study considerably broadens the range of options to this end. Whilst it is possible to do this straightforwardly (e.g. one ester for an overview of the cell), the variety of compartment-specific weighting of nondescript stains needs to be considered carefully. The experiments presented in this paper were conducted on one of the most widely studied human cell lines, HeLa, allowing us to directly compare dye specificities, staining patterns and spatial organisation of compartments. It is our recommendation that an equivalent series of validation is performed when using these dye Esters on different, particularly non-mammalian sample and cell types.

In addition to labelling with dye esters, alternative biomolecules can be implemented to investigate cellular structure, for instance palmitate and maleimides shown in pan-ExM offer another way to investigate the proteome (M’Saad & Bewersdorf, 2020). The distribution of other types of biomolecules has been observed using a labelling strategy termed “fluorescent labelling of abundant reactive entities” (FLARE), whereby hydrazide dyes covalently bind to oxidised carbohydrates, which can be performed in parallel to proteome and DNA labels (Lee et al., 2022; Mao et al., 2020). Furthermore, click chemistry allows the labelling of other biomolecules entirely, such as sphingolipids (Götz et al., 2020).

### Preferential compartment labelling

Based on our characterisation, the compartment-specific weighting of the labelling densities of the dye esters, individually, is determined primarily by the hydrophobicity of each species (see Table 1). In particular, hydrophobic species like the BODIPY NHS esters are generally excluded from the nucleoplasm, but may stain densely in the rough ER networks rich in integral membrane proteins with abundant hydrophilic domains. It is also likely that such exclusion from the nucleoplasm may be absent in human cell types with higher lipid metabolisms (e.g. hepatocytes) that feature nucleoplasmic lipid droplets (Layerenza et al., 2013). By contrast, we observed nucleoplasmic staining with both highly and weakly charged dye esters, suggesting that the overall dye ester charge, if at all, bears only a minor effect on compartment specificity.

Many of the staining patterns and densities that we observe are also composite images of multiple compartment types, particularly as revealed in Fig 3. In our observations, the fluorescent NHS ester are best used for either a.) identifying and distinguishing aqueous spaces that represent lumina or free cytosolic spaces between compartments (e.g. Fig 3A), b.) delineating the overall geometry of large compartments like the ER network as a counter-stain to highly localised or heterogeneous antibody epitopes (e.g. Fig 3B-i), or c.) demonstrating spatial relationships of an antibody epitope or highly localised organelle to neighbouring cellular compartments. Outside of these common experimental scenarios, we have also shown the capacity to visualise the full extent of membraneless compartments such as perinuclear germ granules in *C. elegans* that are formed through liquid-liquid phase separation (Suen et al., 2022).

### The grammar of fluorescent dye ester stainings

A highly promising application of the fluorescent NHS ester staining is the rapid application of the probe and multi-spectral ExM either with multiple dye ester species (Sim et al., 2022) or in conjunction with immunofluorescence (M’Saad & Bewersdorf, 2020). Our findings confirm the modulation of staining intensities of ester species such as TFP AZ488 in a compartment-specific manner in the presence of a simultaneously-applied partner ester species. Conversely, sequential application of dye ester species may also shift the relative densities of staining for a given species depending on the order in which it is applied. The compartment specificity may be more heavily influenced in ExM experiments depending on the application at pre-gelation, inter-digestion and post-digestion stages. These variations may be viewed as additional degrees of versatility of the fluorescent ester probes. We argue that, at the very least, this variation warrants a series validations for each protocol depending on the cell type and timing of the staining steps.

The differences in patterns arising from the exact timepoint of labelling likely depends on 1) the availability of amine groups for esters, and 2) the fluorescence loss of existing ester dyes upon exposure to the free radical gel polymerisation and digestion steps. Availability of amines is likely to be substantially altered following the anchoring step with AcX, which labels amines with methacryloyl groups (allowing their incorporation into the gel matrix). When applied pre-gelation, esters likely have access to more amines (and may in turn lessen the amount that can be anchored), however these esters are then affected by the free radical polymerisation and digestion steps, which will alter brightness and distribution in a heterogeneous manner across the sample. When applied inter-digestion, esters presumably have less of the original amines to bind to (after anchoring), but the digestion process may expose new amines in previously inaccessible regions, and in addition these esters are exposed less to the free radical process following gelation. Esters applied post-digestion are likely binding to leftover amines following the effects of proteinase, and are not subjected to the brightness loss from free radicals or digestion. These differential labelling outcomes emphasise the importance of optimising experiments for each ester, especially determining the best timing of addition. For instance, dyes that are less resistant to bleaching during the expansion process could yield better results when applied post-digestion.

## Conclusions

We have characterised a collection of 14 fluorescent dye esters in HeLa cells, and observed differential compartment labelling with increased resolution from expansion microscopy. By dual-labelling with antibodies we have demonstrated compartment-specificity of the weighting of dye ester labelling densities and how this may be directed to specific compartments depending on the timing of application in ExM. We also report inter-ester interactions which adds complexity to the labelling patterns possible.

## Acknowledgements

The authors declare no conflict of interests.

The NHS ester staining patterns and sub-cellular specificities described in this paper relate to the HeLa cell line. Henrietta Lacks, and the HeLa cell line that was established from her tumor cells without her knowledge or consent in 1951, have made significant contributions to scientific progress and advances in human health. We are grateful to Henrietta Lacks, now deceased, and to her surviving family members for their contributions to biomedical research.

The authors acknowledge funding from the UK Research and Innovation (MR/S03241X/1) and the Integrated Biological Imaging Network (RE13780) pump-priming scheme funded by the Medical Research Council and managed by King’s College London. Imaging work was performed at the Wolfson Light Microscopy Facility at the University of Sheffield.

## Author Contributions

TMDS, KMS, IJ designed research and acquired the funding required. MS, TS, RSS provided materials towards experiments. TMDS performed experiments, made primary observations, carried out data curation, and analysis. IJ provided the supervision. TMDS and IJ wrote the manuscript.

## Methods

### Resource Availability

#### Lead contact

Further information and requests for resources should be directed to and will be fulfilled by the lead contacts, Izzy Jayasinghe or Tom Sheard.

#### Materials Availability

This study did not generate new unique reagents.

#### Data and code availability

Data reported in this paper will be shared by the lead contacts upon request. This paper does not report original code. Any additional information required to reanalyse the data reported in this paper is available from the lead contacts upon request.

### Experimental model and subject details

#### Cell line

HeLa-CCL2 cells (human cervix epitheloid carcinoma), originally sourced from the European Collection of Authenticated Cell Cultures (ECACC 93021013), were gifted to us by the Department of Infection, Immunity and Cardiovascular Disease, Medical School, University of Sheffield, Beech Hill Rd, Sheffield S10 2RX.

Cells were maintained in Dulbecco’s Modified Eagle Medium (Thermo Fisher Scientific) (containing 10% (v/v) foetal bovine serum (Thermo Fisher Scientific) and 1% (v/v) penicillin-streptomycin (Thermo Fisher Scientific)) and were stored in an incubator set at 37 °C and 5% CO2. Cells used in these experiments were from passage numbers between 13 and 25.

Following a passage the cells were counted with a haemocytometer and made into a 75,000 cells/ml solution. 2 ml of this solution (150,000 cells total) was plated onto 22 x 22 mm glass coverslips (Menzel Gläser, thickness #1.5) in the bottom of 6-well plates, which had been coated with 0.01 mg/ml poly-D-lysine (Cultrex). Two days of culture on coverslips enabled the cells to attach and present an elongated morphology, which was preferable to observe structures in the cytoplasm.

Cells on coverslips were fixed at Day 2 by immersion in 2% (v/v) paraformaldehyde (Sigma-Aldrich) made up in phosphate-buffered saline (PBS, Sigma-Aldrich) for 10 minutes at room temperature. Samples were washed three times with PBS for 10 minutes each. Fixed samples were stored until labelling experiments in storage solution (containing 0.05% (w/v) bovine serum albumin (Thermo Fisher Scientific), 0.1% (v/v) sodium azide (Sigma-Aldrich), made up in PBS) in 4°C.

### Fluorescent dye esters

The fluorescent dye esters used have been summarised in Table 1, including details regarding their excitation and emission wavelengths, reported hydrophobicity and charge. The LogD value for each fluorescent ester species was estimated as a measure of its hydrophobicity, using Chemaxon Calculator Plugin (address: https://disco.chemaxon.com/calculators/demo/plugins/logd/).

Ester stock solutions were prepared by reconstituting in DMSO (Biotium) to 10 mg/ml concentration. Stock aliquots were stored in the −20 °C freezer, within a desiccator chamber containing silica. Working solutions of dye esters were prepared by diluting the stock solution to working concentration of 10 μg/ml in ester stain solution, containing 100 mM sodium bicarbonate (Sigma-Aldrich), 1 M sodium chloride (Sigma-Aldrich), made to pH 6 in dH20.

Ester labelling was performed was performed for 1 hour 30 minutes at room temperature, at various stages of the ExM protocol; either pre-gelation, inter-digestion (whereby gels were digested for 4 hours, then incubated with the dye esters, before another 4 hours of digestion), or post-digestion.

### Immunolabelling

Fixed samples were permeabilised with 0.1% (v/v) Triton X-100 (Sigma-Aldrich) in PBS for 10 minutes at room temperature and underwent a blocking step with 0.05% (v/v) Triton X-100 and 10% (v/v) normal goat serum (Thermo Fisher Scientific) in PBS, for one hour at room temperature.

For immunolabelled samples, after blocking the samples were incubated with primary antibodies overnight at 4°C. The primary antibodies used in this study were rabbit polyclonal KDEL (Thermo Fisher Scientific, code: PA1-013), rabbit monoclonal GM130 (abcam, code: ab52649) and mouse monoclonal ATP5A1 (Thermo Fisher Scientific, code: 43-9800). Antibodies were diluted 1:200 in antibody incubation solution (containing 2% (w/v) bovine serum albumin, 0.05% (v/v) Triton X-100, 2% (v/v) normal goat serum, 0.05% (v/v) sodium azide, made up in PBS). The following day, samples were washed three times with PBS for 20 minutes each, and then incubated with secondary antibodies at room temperature for 2 hours. The secondary antibodies used were Alexa Fluor 488 (anti-mouse and anti-rabbit IgG), Alexa Fluor 594 (anti-mouse and anti-rabbit IgG), and Atto647N (anti-rabbit-IgG) diluted 1:200 in antibody incubation solution.

Immunolabelled samples were checked on the microscope to ensure suitable labelling quality and density prior to proceeding with the ExM protocol. Imaging is described in the section Image Acquisition. Sample coverslips were attached to acrylic slides to facilitate viewing on the microscope stage.

### Expansion microscopy

Immunolabelled samples were incubated with 0.1 mg/ml acryloyl-X (Thermo Fisher Scientific) overnight at 4°C for the anchoring step, then washed three times in PBS prior to the addition of gel solution.

4x expanding gels were prepared according to the recipe from the protein retention ExM approach (Tillberg et al., 2016). In short, monomer solution (containing 8.6% (w/v) sodium acrylate (Sigma-Aldrich), 2.5 % (w/v) acrylamide (Sigma-Aldrich), 0.15% (w/v) N,N’-Methylenebisacrylamide (Sigma-Aldrich), 11.7% (w/v) NaCl, PBS) was pre-made and stored in aliquots at −20°C. Samples were incubated in monomer for 30 minutes at 4°C, and was removed before adding the polymerisation solution, made by mixing monomer solution with 0.2% (w/v) ammonium persulfate (Sigma-Aldrich), 0.2% (v/v) N,N,N’,N’-Tetramethylethylenediamine (Sigma-Aldrich), and PBS. This solution was placed onto a parafilm-coated slide, between two coverslip spacers defining the dimensions of the final gel. The coverslip bearing the sample was then inverted onto the blob of gel solution. Polymerisation was enabled at 37°C for two hours.

Polymerised gels were removed from the coverslip chamber, cut into an asymmetric shape (to allow the correct orientation of the gel to be confirmed), and measured to obtain the pre-expansion size. Gels were transferred to a foil-coated 6-well plate for the digestion step, 8 U/ml proteinase K (New England Biolabs) diluted in digestion buffer (containing 50 mM Tris pH 8.0 (Invitrogen), 1 mM ethylenediaminetetraacetic acid (Sigma-Aldrich), 0.5% (v/v) Triton X100, 0.8M guanidine HCl (Sigma-Aldrich) and deionised water (dH_2_0)). Digestion was performed overnight at room temperature, and was stopped early the next morning by removing the digestion solution and adding dH20.

Gels were transferred to foil-coated Petri dishes for the expansion step, which was achieved by five 30-minute washes in dH20. Once expansion had reached a plateau, gels were measured to obtain the post-expansion size.

Prior to imaging, squares of gel were cut and loaded into imaging chambers (formed of a glass coverslip attached to an acrylic slide with a 18×18 mm cut-out). The imaging chamber coverslip was coated with 0.1% (v/v) poly-L-lysine (Sigma-Aldrich) for 30 minutes at room temperature, before three washes in dH20.

### Image acquisition

Airyscan imaging was performed on an inverted LSM 880 Airyscan microscope (Carl Zeiss, Jena), using a 40x oil immersion 1.3 NA objective (working distance 210 μm). Airyscan imaging enables a roughly two-fold improvement in resolution over confocal imaging. Dyes were excited using the following lasers; Argon 488 nm, DPSS 561 nm and HeNe 633 nm, while emission bands were selected using the spectral detector and recorded with the 32-element GaAsP detector. Pixel sampling was 40 nm/pixel.

### Image analysis

Airyscan processing, entailing pixel-reassignment and deconvolution, was performed using the Zen software. Applications of colour-tables and composite multi-channel overlays were performed in FIJI (ImageJ 1.53c). Scalebars presented on images indicate the ‘pre-expansion’ size, as they have been normalised for the expansion factor calculated from the physical gel size. The grey colour-tables of Airyscan and single-colour 4x EExM images presented in the Figures 1 and 2 have been inverted to bring the morphological appearances in line with that of thin-section EM. Therefore, the regions shown in darker pixels represented higher density of staining, therefore fluorescence intensity, compared to pixels that were lighter or white.

**Supplementary figure S1.**
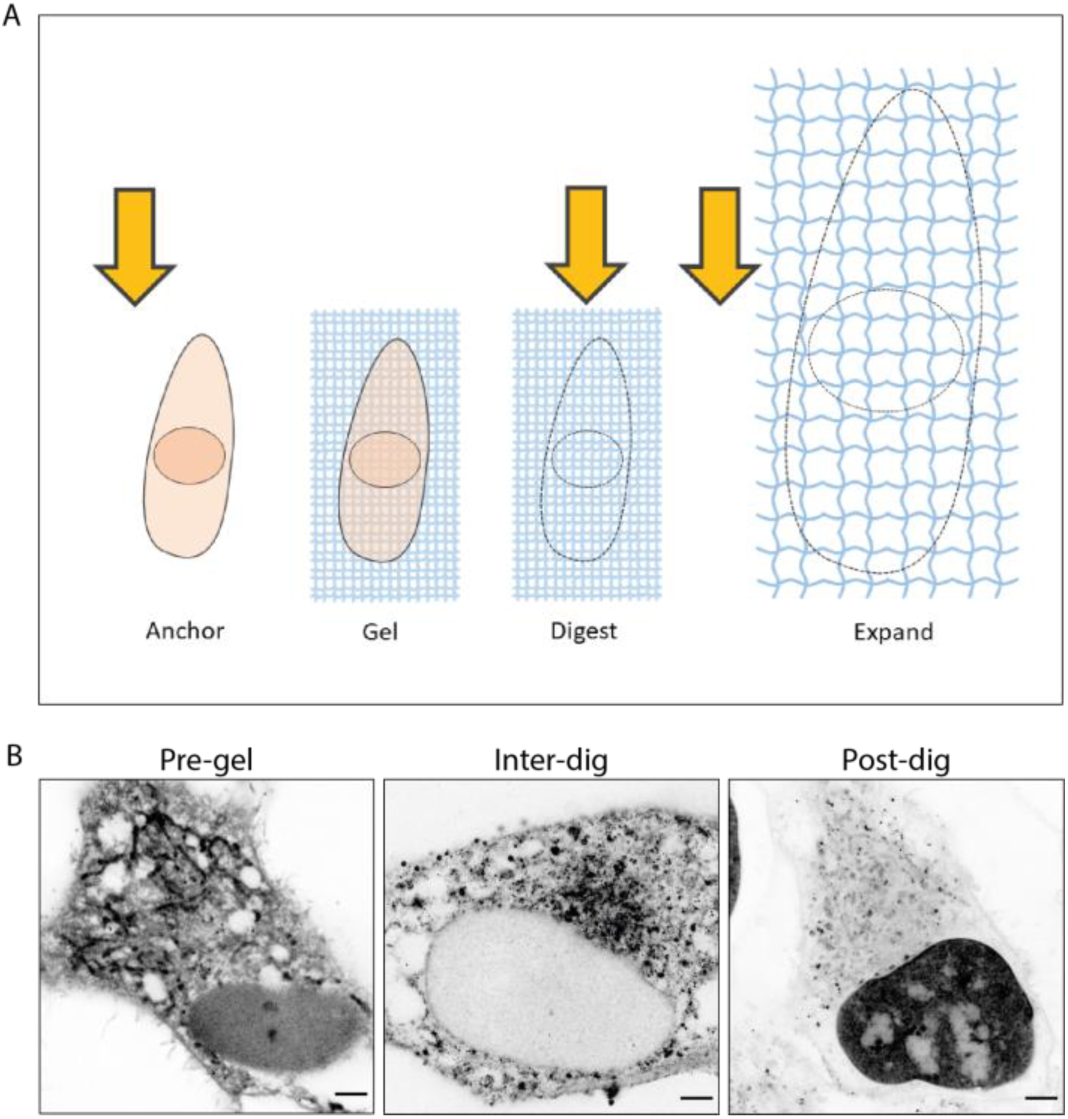
**(A)** Schematic showing the generalised ExM pipeline. Labelling can be performed at different timepoints (highlighted by yellow arrows) either pre-gelation, inter-digestion (4 hours of digestion, ester labelling, another 4 hours digestion), or post-digestion. **(B)** 4x EExM images of NHS ATTO647N applied either pre-gelation, inter-digestion (4 hours of digestion, ester labelling, another 4 hours of digestion), or post-digestion, demonstrate the variety of intracellular structures which can be targeted. Scale bars (expansion factor rescaled): 2.5 μm.

